# Revelation of the Genetic Basis for Convergent Innovative Anal Fin Pigmentation Patterns in Cichlid Fishes

**DOI:** 10.1101/165217

**Authors:** Langyu Gu, Canwei Xia

**Affiliations:** Zoological Institute, University of Basel, Vesalgasse 1, 4051, Basel, Switzerland.; Key Laboratory of Freshwater Fish Reproduction and Development, Ministry of Education, Laboratory of Aquatic Science of Chongqing, School of Life Sciences, 400715, Southwest University, Chongqing, China.; Ministry of Education Key Laboratory for Biodiversity and Ecological Engineering, College of Life Sciences, Beijing Normal University, Beijing, China.

**Keywords:** convergent evolution, novelty, gene network, egg-spots, the blotch, cichlid fishes

## Abstract

Determining whether convergent novelties share a common genetic basis is vital to understanding the extent to which evolution is predictable. The convergent evolution of innovative anal fin pigmentation patterns in cichlid fishes is an ideal model for studying this question. Here, we focused on two patterns: 1) egg-spots, circular pigmentation patterns with different numbers, sizes and positions; and 2) the blotch, irregular pattern with no variation among species. How these two novelties originate and evolve remains unclear. Based on a thorough comparative transcriptomic and genomic analysis, we observed a common genetic basis with high evolutionary rates and similar expression levels between egg-spots and the blotch. But the associations of common genes with transcription factors and signalling pathways in the core gene network, as well as the integration of advantageous genes were observed specifically for egg-spots. Focusing on the whole transcriptomic level instead of limited numbers of candidate genes can explain the opposite conclusion between our study and the one from Santos et al. (2016), which suggested that no common genetic basis is shared between the blotch and egg-spots. Here, we propose that the re-use of the common genetic basis indicates important conservative functions (e.g., toolkit genes) for the origin of these convergent novel phenotypes, whereas independently evolved associations of common genes with transcription factors and signalling pathways (intrinsic factor) can free the evolution of egg-spots, and together with the integration of advantageous genes (extrinsic factor) can provide a clue to link egg-spots as a key innovation to the adaptive radiation in cichlid fishes. This hypothesis will further illuminate the mechanism of the origin and evolution of novelties in a broad sense.

## Introduction

How evolutionary novelties evolve remains an open question in evolutionary biology (Annona et al. 2015; Clark-Hachtel & Tomoyasu 2016; Soltis & Soltis 2016; Yong & Yu 2016). Such novelties provide the raw materials for downstream selection, thereby contributing to biological diversification (Pigliucci & Müller 2010). Examples of evolutionary novelties are neural crest cells in vertebrates (Shimeld & Holland 2000), the beaks of birds (Bhullar et al. 2015; Bright et al. 2016), and eyespots on the wings of nymphalid butterflies (Monteiro 2015). Evolutionary novelty is generally defined in the context of character homology, e.g., as “a *structure that is neither homologous to any structure in the ancestral species nor serially homologous to any part of the same organism*” (Müller & Wagner 1991). However, the inference of homology is not always straightforward (Bang et al. 2002; Cracraft 2005; Panchen 2007; Hall 2013; Faunes et al. 2015). Wagner (Wagner 2007) thus proposed that homology should be inferred using information from the gene network underlying a trait. In this case, characters would be homologous if they shared the same underlying core gene network, whereas novelty would involve the evolution of a quasi-independent gene network, which integrates signals (e.g., input signals and effectors) into a gene expression pattern unique to that character (Wagner 2014). To what extend a gene network is “innovative” compared to the ancestral gene network, and how this gene network affects the evolution of a trait are interesting questions that remain unanswered.

In addition, similar traits can independently evolve in two or more lineages, such as the convergent evolution of echolocation in bats and whales (Shen et al. 2012) and anal fin pigmentation patterns in cichlid fishes (Salzburger et al. 2007; M Emília Santos et al. 2016). Whether convergent phenotypes result from the same gene (e.g., *Mc1r* in mammals (Hoekstra et al. 2006) and birds (San-Jose et al. 2015)) or not (e.g., anti-freezing protein genes (Chen et al. 1997) in Antarctic notothenioid and Arctic cod)), and the extent to which evolution is predictable (Stern 2013) have long been discussed, but no agreement has been reached (Arendt et al. 2008; Stern 2013). Actually, convergent phenotypes can partially share a common genetic basis, for example, independently re-deploy conserved toolkit genes (wing patterns in Heliconius butterfly (Joron et al. 2006) and the repeated emergence of yeasts (Nagy et al. 2014), or different genes within the same pathway (Berens et al. 2015)). Alternatively, these phenotypes can also be derived from deep homology based on ancestral structure (Shubin et al. 2009). Therefore, instead of simply focusing on whether convergent phenotypes are derived from the same genes, it is important to answer the following questions: 1) To what extent does the common genetic basis apply to convergent evolution? 2) Among the genes expressed in independently evolved traits, which genes are conserved? 3) What are the roles of these genes in the evolution of convergent phenotypes? 4) How did these genes evolve in independent lineages, i.e., are they deeply homologized or independently recruited? The answers to these questions at both the individual gene and gene network levels will give a clue about the genetic basis of the origin and evolution of novelty.

East African cichlid fishes, exposed to explosive radiation within millions and even hundreds of thousands of years, are classical evolutionary model species (Kocher 2004; Salzburger 2009). The relatively close genetic background of these fishes and availability of genomics data provide powerful tools to study the relationship between phenotype and genotype (Salzburger 2009; Brawand et al. 2014). Many convergent phenotypes have been identified in cichlid fishes, such as thick lip (Colombo et al. 2013), lower pharyngeal jaw (Muschick et al. 2012), and the convergent evolution of anal fin pigmentation patterns, a new model to study the origin of evolutionary novelty (Salzburger et al. 2007). Convergent evolution of anal fin pigmentation patterns primarily include two patterns: egg-spots, i.e. conspicuous pigmentation patterns with a circular boundary, originate once in the most species-rich lineage, i.e. haplochromine lineage (Salzburger et al. 2005a, 2007); and the blotch, with an irregular boundary that only present in four subspecies within *Callochromis* genus in the ectodine lineage (Brichard 1989) (Figure 1). Egg-spots exhibit large varieties (different numbers, sizes and positions, etc.) both within and among different species (Santos & Salzburger 2012; Theis et al. 2017). Both egg-spots and the blotch have been associated with sexual selection (Wickler 1962; Hert 1989; Theis et al. 2012, 2015).

**Figure 1.**
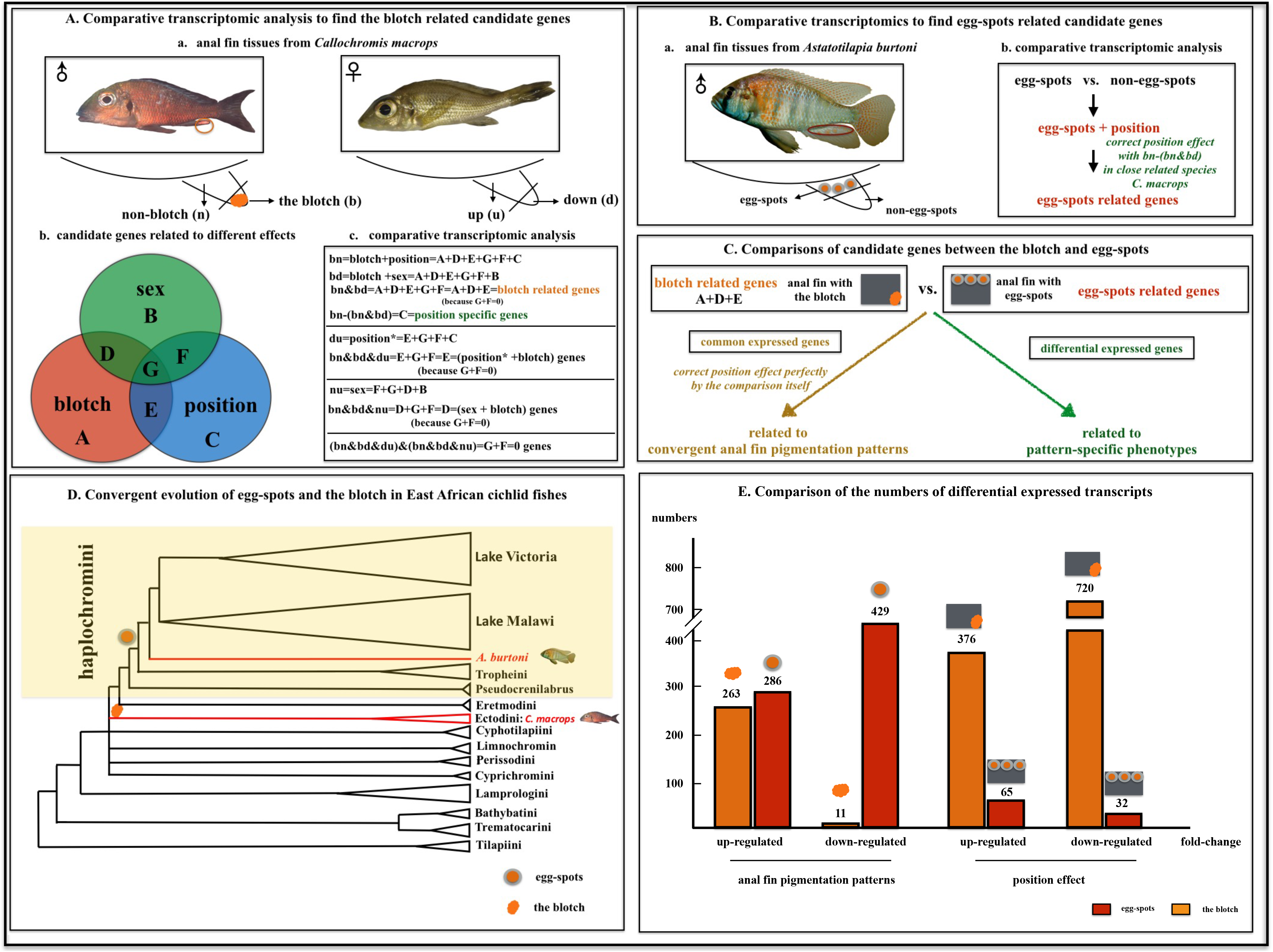
Comparative transcriptomic analysis to find candidate genes for the blotch and egg-spots. (A) a. Four kinds of tissues (blotch (b) and non-blotch (n) in males; corresponding position in females, up (u) and down (d)). b. Different effects can be involved in the candidate genes resulting from different comparisons. c. Different comparisons to find candidate genes for the blotch. (B) a. Two kinds of tissues (egg-spots and non-egg-spots) were used in the males of *A. burtoni*, and since the females also possess egg-spots, they cannot be used as the control. b. Comparative transcriptomic analyses to find candidate genes of egg-spots with position effect controlling. (C) Comparisons to find common and pattern specific candidates for the blotch and egg-spots. (D) Convergent evolution of egg-spots (originating once in the most species-rich lineage, haplochromine lineage) and the blotch (ectodine lineage). The area of the triangle represents the species richness based on the study from (Salzburger et al. 2005b) and <u>https://en.wikipedia.org</u>. The schematic molecular phylogeny was based on the combined evidences from previous studies (Salzburger et al. 2005b; Meyer et al. 2015; Takahashi & Sota 2016). (E) The numbers of candidate transcripts.

By far, only a few studies begun to dissect the genetic basis of these anal fin pigmentation patterns. The first one came from Salzburger et al. (Salzburger et al. 2007), which suggested that a xanthophore related gene, *csf1ra*, is expressed in egg-spots and has undergone adaptive sequence evolution. Subsequently, Santos et al. (Santos et al. 2014) found a cis-regulatory element insertion in the upstream region of *fhl2b* segregating with egg-spots can drive the expression of green fluorescent protein (GFP) in iridophore in zebrafish, and could be causally linked to the origin of egg-spots. More recently, Santos et al. (2016) partially touched on the genetic basis of the blotch by detecting the gene expression profiles of 46 out of 1229 egg-spots related candidate genes and found few common expression patterns between the blotch and egg-spots; thus, concluded that these two traits do not have the same genetic basis. However, this inference was only based on the results from a small portion of candidate genes (3.7%); and the investigated genes were top egg-spots specific candidate genes that are unlikely to reflect the expression pattern of the blotch. Therefore, it is still too early to draw this conclusion.

To investigate the genetic basis of the origin and evolution of these two convergent novelties, we conducted a thorough comparative transcriptomic and genomic analysis with respect to a gene network, and proposed the origin and evolution of these two convergent novel phenotypes in an evolutionary scope. Since these two novelties are both pigmentation patterns with similar colours, albeit with different patterns, we propose that they share a common genetic basis, but independent associations with different genes and pathways can be responsible for their differences.

## Results

### Differential expressed (DE) transcripts for the blotch and egg-spots

Illumina-based RNA sequencing of anal fin tissues of *Callochromis macrops* generated between 15 to 23 million raw reads per library with average read quality 28. After adaptor trimming, between eight to ten million reads per library were finally mapped to the Tilapia transcriptome assembly available from the Broad Institute (ftp://ftp.ensembl.org/pub/release-81/fasta/oreochromis_niloticus/cdna/. version 0.71) (Additional File 1). Illumina reads are available from the Sequence Read Archive (SRA) at NCBI under accession number SRA SRP082469. After position and sexual effects controlling, we identified 263 up-regulated DE transcripts and 11 down-regulated DE transcripts for the blotch in *C. macrops*; 286 up-regulated DE transcripts and 429 down-regulated DE transcripts for egg-spots in *Astatotilapia burtoni* (Figure 1E).

### Comparative transcriptomic analyses for the blotch and egg-spots

Forty-four transcripts showed similar expression patterns in both the blotch and egg-spots. Particularly for the top ten blotch DE transcripts, six showed similar expression patterns in egg-spots, corresponding to two duplicated iridophore-related genes *(pnp4a* and *pnp5a*), the immunity-related gene *ly6d*, the ligand transporter-related gene *(ApoD)*, and the two genes associated with egg-spots formation *(fhl2a* and *fhl2b)* (Santos et al. 2014) (Table 1). Besides, two transcripts were highly expressed in egg-spots, but low expressed in the blotch, corresponding to a novel transcript ENSONIT00000023612 and the immunity-related gene *steap4* (Benard et al. 2014) (Table 2). Fifteen transcripts were over-expressed in the blotch but down-regulated in egg-spots (Table 2).

**Table 1.**
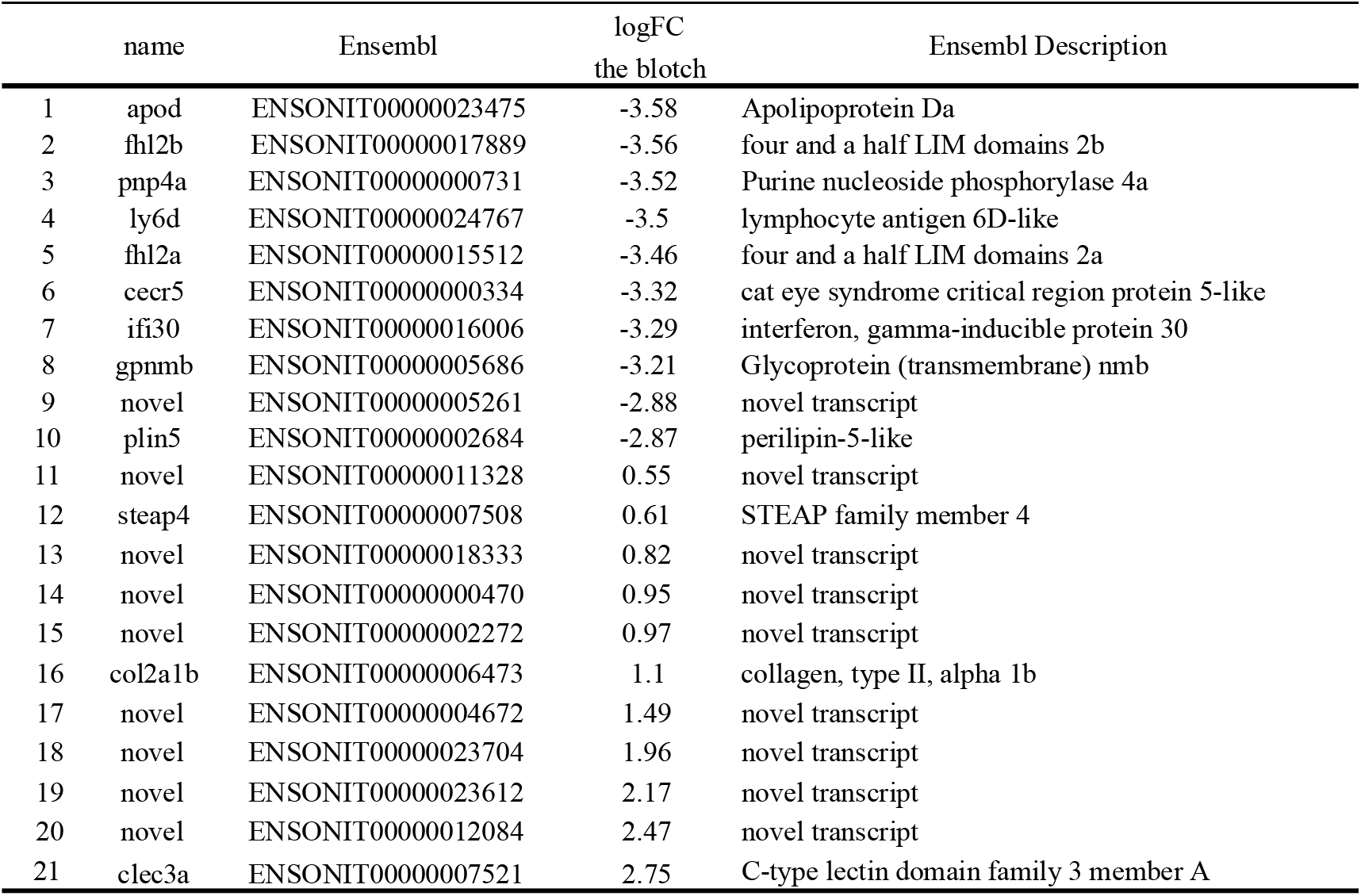
Top ten common blotch DE transcripts and 11 down-regulated blotch DE transcripts

**Table 2.**
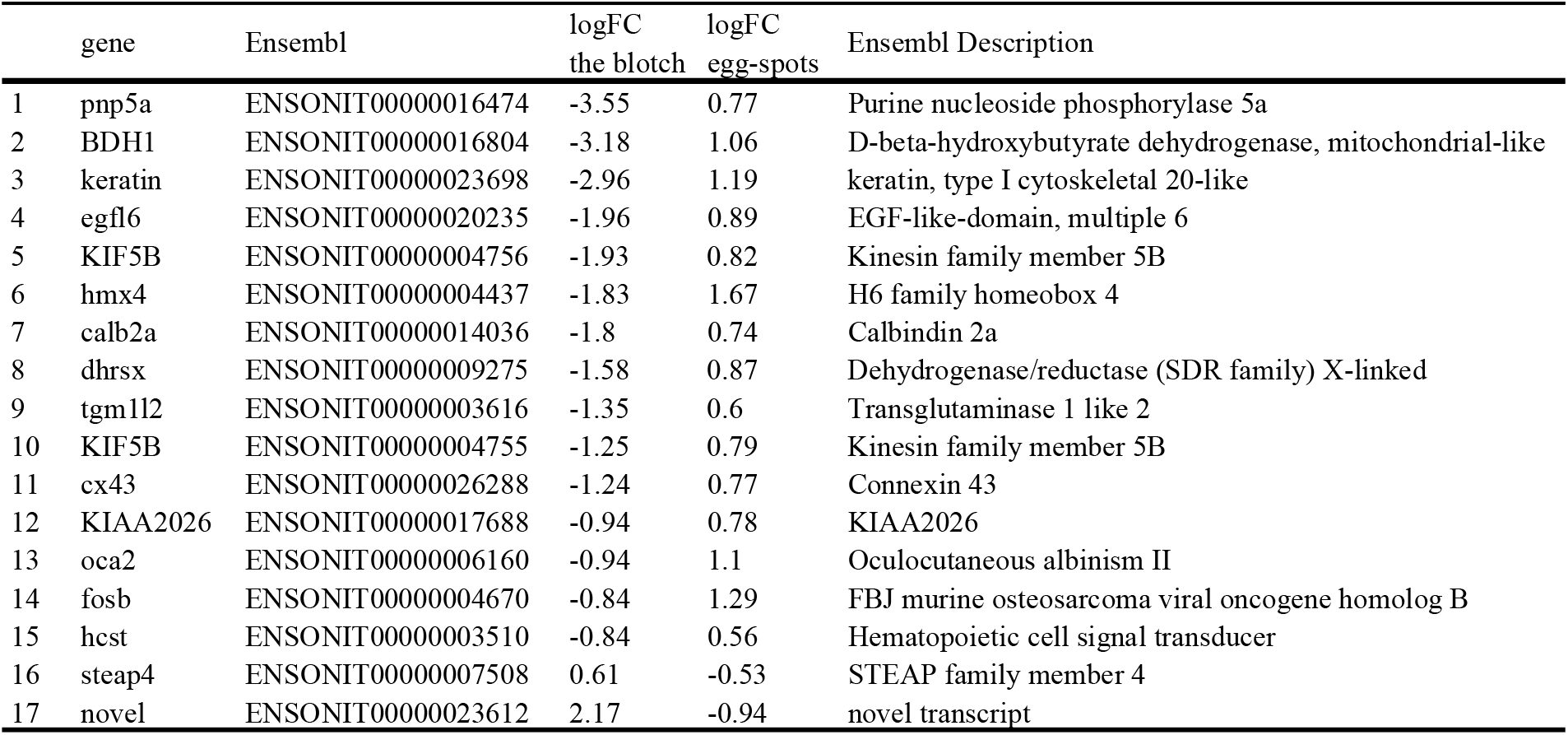
Differential expressed transcripts with contrasting expression patterns between the blotch and egg-spots

Clustering analysis based on the expression levels of 44 common transcripts clustered anal fin tissues possessing egg-spots and the blotch together, despite their species boundary (Figure 2B). This was not the case for pattern specific DE transcripts (Figure 2A and 2C). Since sexual effect has already been controlled (Figure 1), sex difference was not the reason here. More than 50% Gene Ontology (GO) terms and pathways were shared between the blotch and egg-spots (Figure 3). Pigmentation related GO terms, including pigmentation cell differentiation (GO: 0050931), pigment granule organization (GO: 0048753), melano-related terms (GO:0030318, GO: 0032438, GO: 0004962), iridophore-related GO term (GO: 0050935), and nucleobase-containing small molecule metabolic process (GO: 0055086) were enriched (Figure 3). Noticeably, there were more egg-spots specific pathways compared to the blotch (Figure 3 and Figure 4).

**Figure 2.**
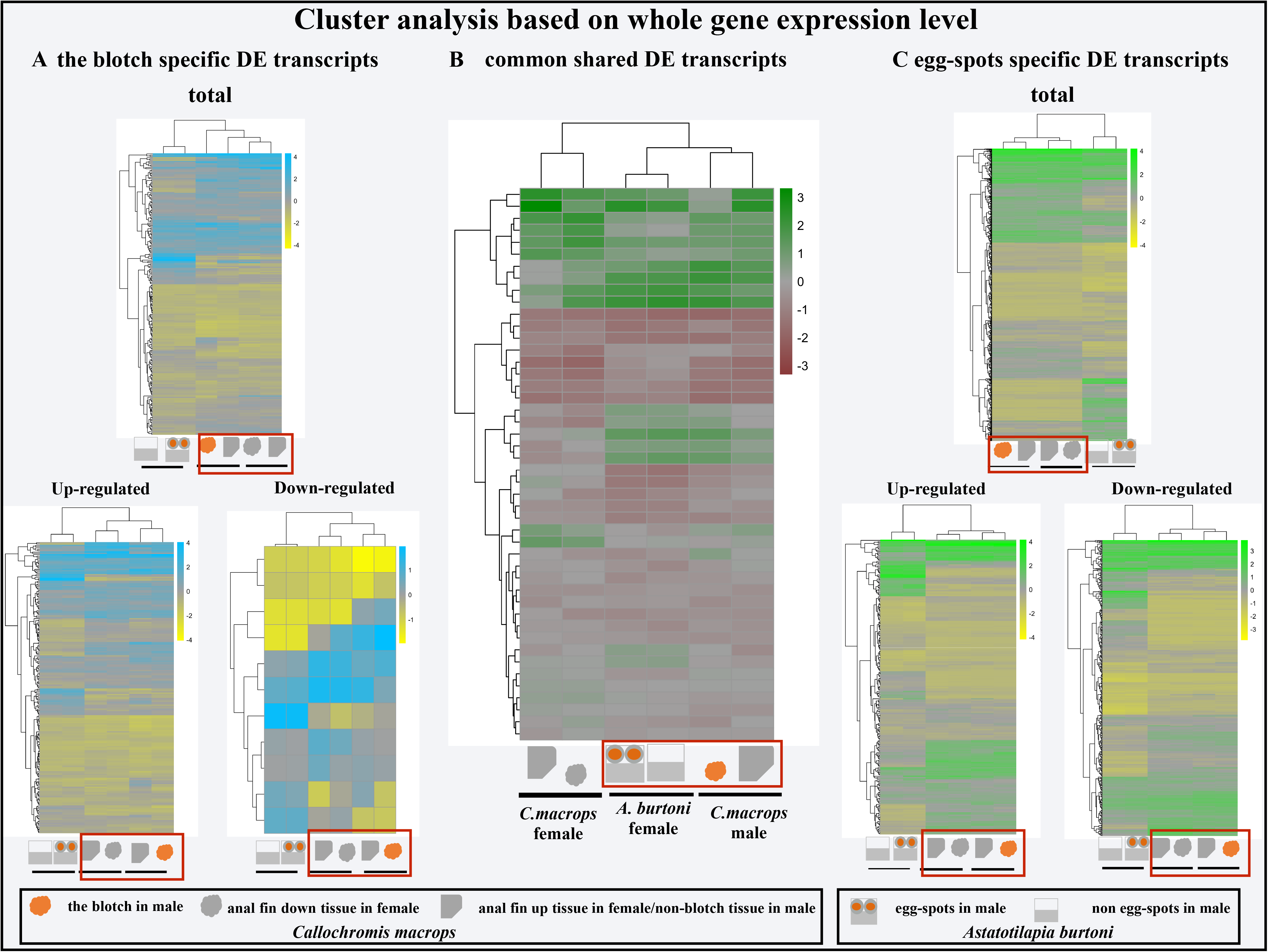
Clustering analysis based on the expression levels of common differentially expressed (DE) transcripts and pattern specific DE transcripts of anal fin pigmentation patterns. The cluster analysis was conducted with the blotch and non-blotch tissues in the male and the corresponding tissues (up tissue and down tissue) in the female of cichlid fish *Callochromis macrops*, and egg-spots and non-egg-spots tissues in the male of cichlid fish *Astatotilapia burtoni.* Orange rectangles showed the clustered anal fin tissues.

**Figure 3.**
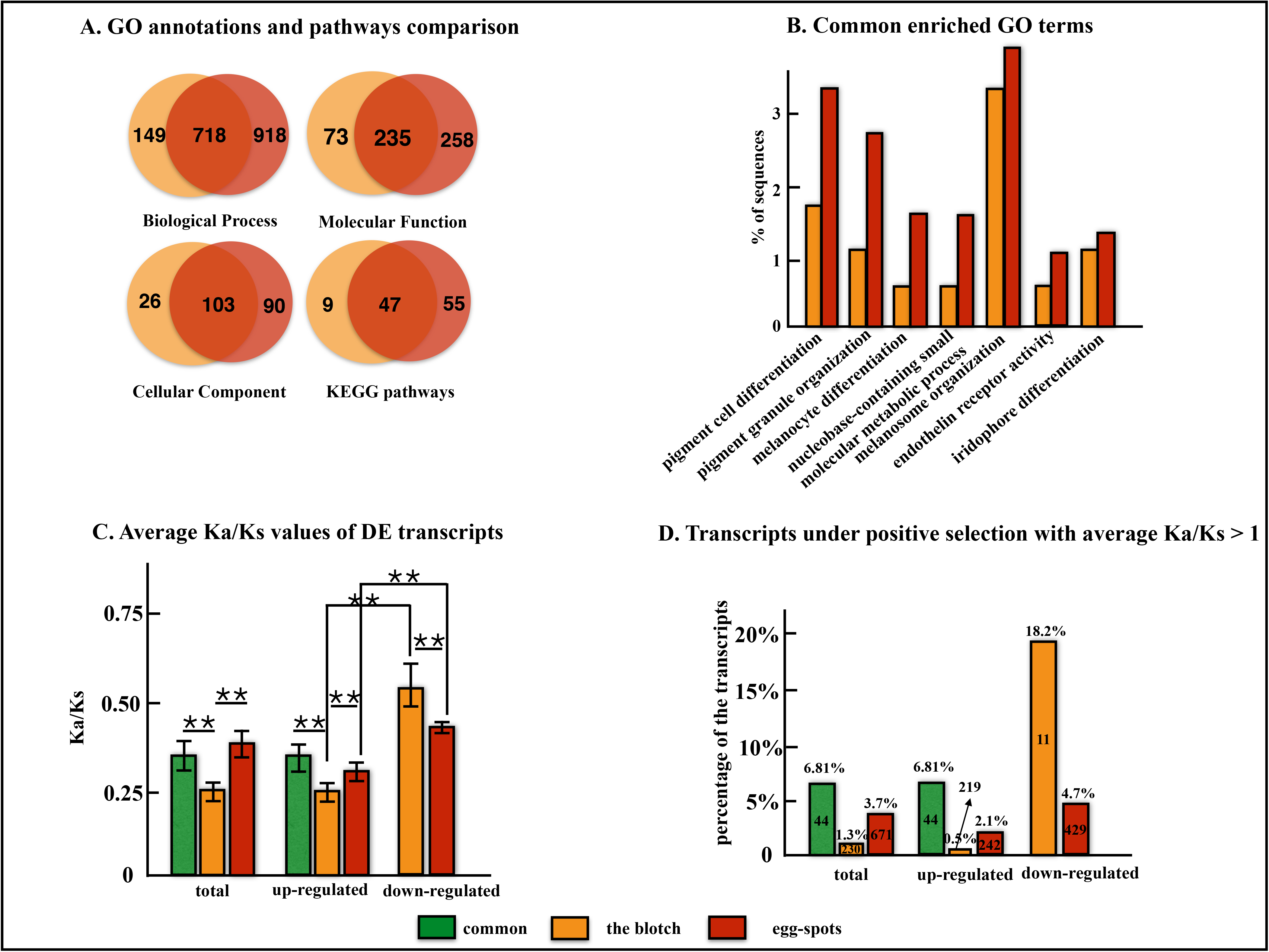
A common genetic basis was shared between the blotch and egg-spots with accelerated evolutionary rates. (A) > 50% common shared gene ontology (GO) terms and pathways between the blotch and egg-spots. (B) Common enriched GO terms between the blotch and egg-spots (FDR<0.1). (C) Average evolutionary rates (Ka/Ks) of common shared differentially expressed (DE) transcripts, the blotch-specific DE transcripts and egg-spots-specific DE transcripts. ** represents p<0.01. (D) Statistics of the transcripts under positive selection with average Ka/Ks values larger than 1. Total numbers were given in the boxes.

**Figure 4.**
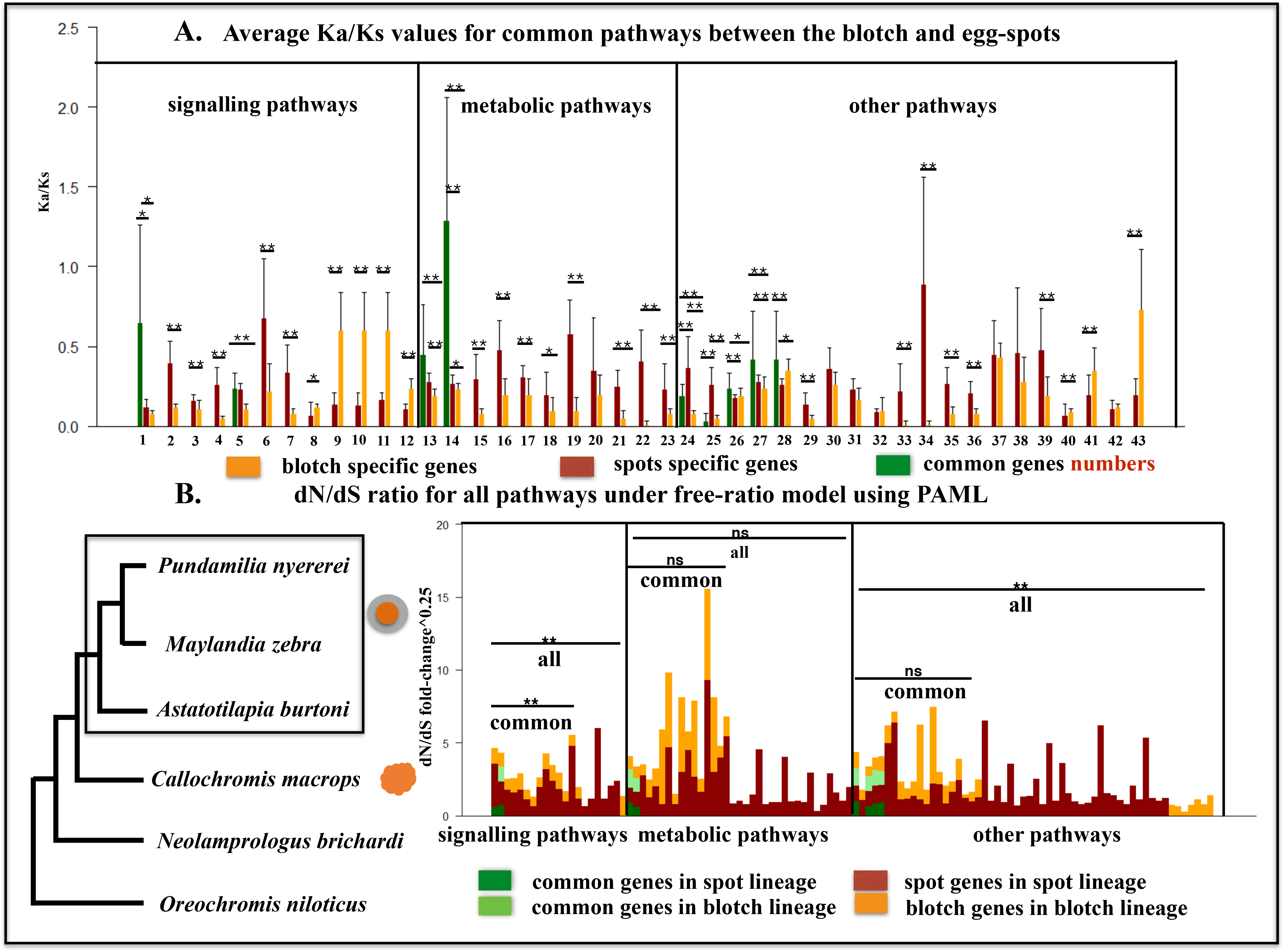
Evolutionary rates calculations for anal fin pigmentation differentially expressed (DE) transcripts related pathways. (A) Average evolutionary rates (Ka/Ks) values of common shared pathways between the blotch and egg-spots. To keep the comparison unbiased, gene pairs with any NA (not applicable or infinite) value occurred were removed (Wang et al. 2011), and thus, 43 common pathways were compared in total. For details see Additional File 2. * represents p<0.05; and ** represents p<0.01. (B) Rates of evolution (dN/dS) for all pathways under free-ratio model in a lineage-specific manner within codeml in PAML. Fold-change ratios of dN/dS in the corresponding lineage were calculated and compared. For details see Additional File 2. **represents p<0.01; ns represents no significance; common represents the common shared pathways; and all represents all the related pathways. Species *Pundamilia nyererei, Maylandia zebra, Astatotilapia burtoni* possess egg-spots, species *Callochromis macrops* possess the blotch, species *Neolamprologus brichardi*, *Oreochromis niloticus* do not possess either the blotch or egg-spots. The phylogeny was based on studies from (Meyer et al. 2015; Brawand et al. 2014; Salzburger et al. 2005a, 2007).

### Evolutionary rate detection for DE transcripts and related pathways

The average Ka/Ks values showed significantly higher rates for common transcripts than blotch transcripts, but not higher than egg-spots transcripts (Figure 3). Common transcripts also possessed the largest percentage of transcripts under positive selection (average Ka/Ks>1) (Figure 3). Interestingly, despite the large difference in the numbers of down-regulated transcripts between the blotch (11) and egg-spots (429), they both showed significantly higher evolutionary rates than up-regulated transcripts (Figure 3). This pattern can also be found for the percentage of transcripts under positive selection (Figure 3). Noticeably, more than half of the transcripts under positive selection are unannotated novel transcripts (Table 3).

**Table 3.** Transcripts under positive selection with average Ka/Ks larger than 1

| Name | ID | Ka/Ks | expression | tissues | Ensembl Description |
| --- | --- | --- | --- | --- | --- |
| map7a | ENSONIT00000000906 | 1.37 | up-regulated | both | Microtubule-associated protein 7a |
| uraha | ENSONIT00000007140 | 1.29 | up-regulated | both | Urate (5-hydroxyiso-) hydrolase a |
| FAM180A | ENSONIT00000013027 | 1.004 | up-regulated | both | Family with sequence similarity 180 member A |
| bscl2l | ENSONIT00000006721 | 1.23 | up-regulated | the blotch | Bernardinelli-Seip congenital lipodystrophy 2, like |
| novel | ENSONIT00000000470 | 1.89 | down-regulated | the blotch | unannotated |
| novel | ENSONIT00000012084 | 1.12 | down-regulated | the blotch | unannotated |
| novel | ENSONIT00000002070 | 1.65 | up-regulated | egg-spots | unannotated |
| novel | ENSONIT00000010127 | 1.14 | up-regulated | egg-spots | unannotated |
| novel | ENSONIT00000022442 | 1.12 | up-regulated | egg-spots | unannotated |
| novel | ENSONIT00000025818 | 1.1 | up-regulated | egg-spots | unannotated |
| spam1 | ENSONIT00000005048 | 1.02 | up-regulated | egg-spots | sperm adhesion molecule 1 |
| novel | ENSONIT00000025871 | 3.61 | down-regulated | egg-spots | unannotated |
| novel | ENSONIT00000026449 | 3.13 | down-regulated | egg-spots | unannotated |
| trappc2 | ENSONIT00000020233 | 2.32 | down-regulated | egg-spots | Trafficking protein particle complex 2 |
| novel | ENSONIT00000016236 | 2.13 | down-regulated | egg-spots | unannotated |
| novel | ENSONIT00000017692 | 1.94 | down-regulated | egg-spots | unannotated |
| novel | ENSONIT00000006770 | 1.66 | down-regulated | egg-spots | unannotated |
| novel | ENSONIT00000000523 | 1.48 | down-regulated | egg-spots | unannotated |
| novel | ENSONIT00000025911 | 1.47 | down-regulated | egg-spots | unannotated |
| mgst3a | ENSONIT00000023478 | 1.4 | down-regulated | egg-spots | Microsomal glutathione S-transferase 3a |
| novel | ENSONIT00000021326 | 1.4 | down-regulated | egg-spots | unannotated |
| novel | ENSONIT00000005252 | 1.15 | down-regulated | egg-spots | unannotated |
| novel | ENSONIT00000004956 | 1.14 | down-regulated | egg-spots | unannotated |
| crema | ENSONIT00000018860 | 1.11 | down-regulated | egg-spots | CAMP responsive element modulator a |
| isg15 | ENSONIT00000026211 | 1.09 | down-regulated | egg-spots | ISG15 ubiquitin-like modifier |
| igfbp6a | ENSONIT00000024154 | 1.08 | down-regulated | egg-spots | Insulin-like growth factor binding protein 6a |
| timp2b | ENSONIT00000006684 | 1.07 | down-regulated | egg-spots | TIMP metalloproteinase inhibitor 2b |
| OLA1 | ENSONIT00000010893 | 1.06 | down-regulated | egg-spots | Obg like ATPase 1 |
| stard14 | ENSONIT00000003484 | 1.06 | down-regulated | egg-spots | START domain containing 14 |
| novel | ENSONIT00000001384 | 1.04 | down-regulated | egg-spots | unannotated |
| novel | ENSONIT00000009645 | 1.03 | down-regulated | egg-spots | unannotated |

To detect the evolutionary rates of related pathways, we calculated average Ka/Ks values for common pathways and dN/dS values under the free-ratio model for all the pathways (Figure 4). Consistent patterns were found using these two different methods. 1) In general, common transcripts showed higher evolutionary rates than the blotch transcripts within the same pathway, confirming our previous results. 2) Egg-spots transcripts showed higher evolutionary rates, primarily in signalling pathways, i.e. 7 out of 12 (58.3%) pathways under average Ka/Ks value calculations (Figure 4a), and were statistically significant under the free-ratio model (common**) (Figure 4b). The three pathways (9,10 and 11) in Figure 4a with high average Ka/Ks values for the blotch were caused by only one unannotated transcript, ENSONIT00000016796. However, under the free-ratio model after lineage effect controlling, the effect of this transcript was minimized (Figure 4b), further confirming the higher evolutionary rates of egg-spots transcripts in signalling pathways. 3) There was no large difference in the evolutionary rates of other pathways between egg-spots and the blotch, i.e. less than half showed higher average rates in either pattern (Figure 4a and 4b), and no statistical significance was detected under the free-ratio model; however, more egg-spots specific pathways, together with common pathways, showed higher evolutionary rates than the blotch (all**) (Figure 4b). Noticeably, different patterns were found for the metabolic pathways in different tests, i.e. higher average Ka/Ks values were found for egg-spots (Figure 4a), but no significance was detected under the free-ratio model (Figure 4b). This finding could be because the later method considers the lineage effect. However, egg-spots possess more specific metabolic pathways than the blotch (Figure 4b, Additional file 2).

### First common neighbour gene networks (FCNs) of the blotch and egg-spots

To characterize the interactions of DE transcripts, we examined their protein-protein interaction networks. Network enrichment analysis in both the blotch and egg-spots showed that the corresponding genes had more interactions among themselves than what would be expected from the genome (p<0.01), suggesting that these genes are at least partially biologically related as a group, further confirming the robustness of our experimental design. To predict the roles of the common genes in the network, we analysed their FCNs (Figure 5), as direct interactions among genes can indicate their roles to the largest extant. These FCNs showed significantly higher interaction degrees than the total gene network (Table 4), reflecting their core roles in the network.

**Figure 5.**
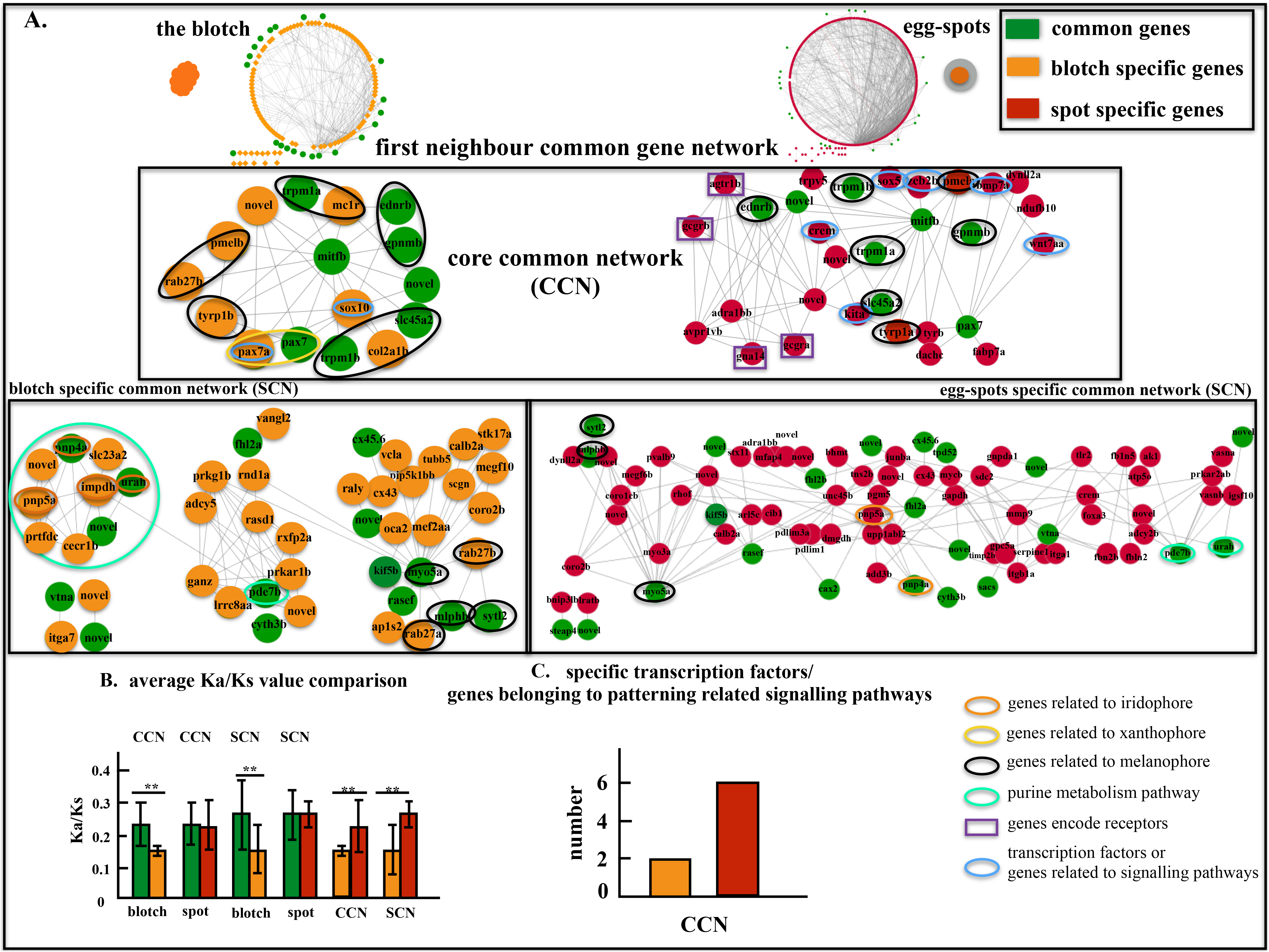
First common neighbour gene interaction network (FCN) analysis for the blotch and egg-spots. (A) FCNs between the blotch and egg-spots, including the core common gene network (CCN) primarily comprising the direct interactions among pigment cells (above), and the specific core gene network (SCN) comprising the interactions among common genes and pattern specific genes (below). (B) Average evolutionary rates (Ka/Ks) comparison of different genes of the CCN and SCN in the blotch and egg-spots. **represents p<0.01. (C) Comparison of the percentage of pattern specific transcription factors (TFs) and genes belonging to the patterning related signalling pathways in CCNs between the blotch and egg-spots.

**Table 4.** Statistics of interaction degree of first common neighbour gene networks

| comparison | average | p-value | significance |
| --- | --- | --- | --- |
| egg-spots fcn vs. egg-spots total | 8.62 vs. 5.66 | 0 | ** |
| egg-spots ccn vs. egg-spots total | 8.74 vs. 5.66 | 0.02 | * |
| egg-spots scn vs. egg-spots total | 8.58 vs. 5.66 | 0 | ** |
| the blotch fcn vs. the blotch total | 6.28 vs. 4.11 | 0 | ** |
| the blotch ccn vs. the blotch total | 6.63 vs. 4.11 | 0.01 | * |
| the blotch scn vs. the blotch total | 6.17 vs. 4.11 | 0 | ** |

fcn, first neighbour gene network; ccn, core common network; scn, specific common network.

Two types of FCNs were formed: 1) direct interactions among the common genes, comprising pigmentation (melanophore and xanthophore)-related genes (core common network, CCN) (Figure 5). Noticeably, unlike in the blotch, the CCN in egg-spots integrated more patterning-related transcription factors (TFs) and genes belonging to patterning-related signalling pathways. 2) The second type of FCN involved interactions among common genes and pattern specific DE genes (specific common network, SCN). Unlike the CCN, the common genes in SCN were only loosely associated (Figure 5). For example, the SCN of the blotch could clearly be divided into sub-networks, connected only by two common genes *(myo5aa* and *pde7b)* (Figure 5). In addition, genes related to purine metabolism pathways were detected in the SCN of the blotch (Figure 5).

The evolutionary rates of common genes were significantly higher than those of pattern specific genes in both CCN and SCN of the blotch, but not those of egg-spots (Figure 5), confirming our previous results (Figure 3). In addition, several novel genes were involved in FCNs (Figure 5). Since the interactions among genes indicate their similar functions, the network presented here enabled the prediction of new gene functions. For example, in the CCN of the blotch, the novel gene ENSONIG00000017631 interacted with *mtifb, mc1r* and *sox10*, suggesting a similar role related to melanin (Hou et al. 2006; San-Jose et al. 2015). The blotch-specific novel gene ENSONIG00000021415 interacted with *mitfb, pax7a* and *pax7b*, suggesting a xanthophore-related role (Minchin & Hughes 2008; Curran et al. 2010).

## Discussion

The thorough comparative transcriptomic and genomic data analysis conducted herein revealed a common genetic basis between the blotch and egg-spots. Focusing on the whole transcriptomic expression level instead of limited numbers of candidate genes can explain the opposite conclusion between our study and the one from Santos et al. (M Emília Santos et al. 2016). Here, we proposed that the common genetic basis is essential for the emergence of these novel phenotypes, and independently evolved associations among common genes with TFs and signalling pathways, as well as the integration of advantageous genes are crucial for the evolution of egg-spots.

### Common genetic basis is important for the origin of the blotch and egg-spots

Similar gene expression patterns in convergent taxa are generally associated with conserved and important roles (such as toolkit genes) in the origin of phenotype (Khaitovich et al. 2006; Pankey et al. 2014). In the present study, clustering analysis based on expression level revealed that convergent evolution of anal fin pigmentation patterns is in parallel with gene expression level, at least on the common genetic basis level. Genes *fhl2a* and *fhl2b*, which were suggested to be associated with egg-spots formation (Santos et al. 2014), were also observed ranking at the top of the blotch candidate gene list, further confirming our hypothesis. Common transcripts showed higher accelerated evolutionary rates in total and in the shared pathways, as well as possessed the highest percentage of transcripts under positive selection, evincing their advantageous roles during evolution. The higher average Ka/Ks value of egg-spots transcripts than the result from Santos et al. (M Emília Santos et al. 2016) could be explained by the integration of recent duplicated genes into our analysis, which is too important to ignore (Glasauer & Neuhauss 2014). High evolutionary rates can be explained by the reasoning that the trait itself is under selection, or alternatively that the formation of the trait integrated advantageous genes in the network, which conserved the phenotype during evolution. Since common transcripts also showed significantly higher evolutionary rates than pattern specific transcripts, we prefer the later scenario. In any case, they both suggest that the common genetic basis is important for the origin of these convergent anal fin pigmentation patterns.

Then what is the common genetic basis and its role in the convergent evolution of anal fin pigmentation patterns? Both GO enrichment and gene network construction showed that the common genes primarily comprise pigmentation-related genes. Indeed, interactions among pigment cells are important for pigmentation pattern formation (Parichy 2009; Parichy & Spiewak 2015; Watanabe & Kondo 2015; Kondo 2017). For example, the mutation of melanophore-related genes *(mitfb, gpnmb, trpm1a, ednrb, trpm1b* and *sox10)*, the iridophore-related gene *(pnp4a*), the xanthophore-related gene *(pax7)* and the connexion-related gene *cx45.6* can induce irregular stripe pattern formation in zebrafish (Shin et al. 1999; Hou et al. 2006; Minchin & Hughes 2008; Curran et al. 2010; Zhang et al. 2012; Reissmann & Ludwig 2013; Irion et al. 2014), suggesting their important roles in the corresponding core gene network. The genes *fhl2b* (iridophore-related gene), *pax7a* (xanthophore-related gene) and *melanocortin* were also reported to be associated with pigmentation pattern formation either on the anal fin *(fhl2b)* or in skin *(pax7a* and *melanocortin)* in cichlid fishes (Salzburger et al. 2007; Santos et al. 2014; Dijkstra et al. 2017). These genes were in the common candidate gene list in the present study, which can have similar roles in the core gene network of anal fin pigmentation pattern formation. This was further confirmed by the finding that the FCNs showed a significantly higher interaction degree than the average degree of total gene network.

In addition, signalling pathways and metabolism pathways occupied a large portion of the shared pathways between the blotch and egg-spots. It is unsurprising that metabolism pathways were involved in these sexually related traits, considering the important trade-off of energy allocation between the survival and reproduction (Bleu et al. 2016). For example, both patterns contain carotenoid-based reddish colours, a limited resource used in the immune system, which in fish, can only be obtained through diet; it is important to trade off its allocation between immune response and sexual signalling (Theis et al. 2017). In addition, many patterning related signalling pathways were shared between the blotch and egg-spots, such as the Wnt signalling pathway, ErbB signalling pathway and MAPK signalling pathway (Budi et al. 2008; Martin et al. 2012; Schneider et al. 2012; Plestant & Anton 2013; Martin & Reed 2014), which can have similar roles here and are advantageous for the emergence of these novel patterns. The re-use of the same large portion of the metabolic and signalling pathways again indicated their importance in the formation of both egg-spots and the blotch.

### Independently evolved associations among common genes and pattern specific genes are crucial for the evolution of egg-spots

Although both the blotch and egg-spots are novelties, their patterns are quite different. The blotch is a reddish pattern with an irregular boundary, exhibiting no varieties among species (Brichard 1989); whereas egg-spots are with conspicuous outer rings and exhibit high evolvability including different numbers, sizes and positions within and among species (Santos et al. 2014; Theis et al. 2017). Egg-spots originate once in haplochromine lineage, the most species-richness lineage, and was suggested to be one of the key innovations associated with the adaptive radiation of cichlid fishes based on phylogeny and character reconstruction (Salzburger et al. 2005a, 2007; Salzburger 2009). It is important to understand the mechanism underlying the different patterns and evolvability between the blotch and egg-spots.

Genes are not isolated from each other, but rather remain connected. Although common genes were shared, more specific pathways were found in egg-spots, especially for signalling pathways and metabolic pathways. Additionally, more direct interactions were observed among common genes for egg-spots genes than those of the blotch in FCNs, especially for the connections with patterning related TFs *(crem, sox5* and *zebΞb)*, and genes belonging to patterning related signalling pathways *(bmp7a, wnt7aa* and *kita)* in the CCN of egg-spots, compared to the only two pigmentation related TFs *(pax7a* and *sox10)* of the blotch (Figure 5) (https://www.ncbi.nlm.nih.gov/gene/). Gene network integrating patterning related TFs and signalling pathways can become relatively independent to integrate signals into a gene expression pattern unique to that character, representing novelty (Wagner 2014), similar to the case of butterfly eyespot (Shirai et al. 2012). The extent of independence of a gene network is important for the evolution of a trait (Wagner 2014): the relatively independent network can free the evolution of egg-spots from the ancestral anal fin network compared to the blotch (intrinsic factor, developmental constraint). In addition, the DE transcripts of egg-spots showed higher evolutionary rates than those of the blotch, both in total and in FCNs, especially in the signalling pathways. Additionally, there are larger percentages of transcripts under positive selection in egg-spots than in the blotch. These findings further suggest that evolutionary advantageous genes can be recruited in the egg-spots gene network, which can be causally linked to the high evolvability of egg-spots (extrinsic factor, selection).

Noticeably, there was a large difference in the numbers of down-regulated DE transcripts between the blotch (11) and egg-spots (429). This difference could be due to the effect of lineage-specific position related transcripts that were not ruled out based on our strategy. However, this explanation is unlikely, considering the very closely related genetic background in cichlid fishes (Baldo et al. 2011; Brawand et al. 2014) and the relatively conservative fin developmental constraint (Maxwell & Wilson 2013; Paço & Freitas 2017). In addition, after position effect controlling, we observed similar numbers of up-regulated DE transcripts between egg-spots and the blotch, suggesting that the comparison strategy was not the issue. Furthermore, down-regulated transcripts of both the blotch and egg-spots showed the same results (clustering analyses and evolutionary rates detection), suggesting that position effect was not the issue, since the blotch transcripts have already controlled position effect.

One explanation could be the importance of down-regulated transcripts in patterning, especially for egg-spots, although this issue is largely ignored. Indeed, down-regulated transcripts showed significantly higher evolutionary rates than up-regulated transcripts in both the blotch and egg-spots, and possessed a large percentage of transcripts under positive selection, suggesting their advantageous roles. Especially, most of which were novel transcripts, reflecting the novelty. Considering the more complicated pattern of egg-spots compared to the blotch, more down-regulated genes could be essential for its formation. Noticeably, unlike the blotch, there was only a small number of position effect genes in egg-spots (Figure 1E), reflecting the relatively homogenous genetic background of anal fin tissue possessing egg-spots in *A. burtoni*, which could also be one of the factors prompting the diversification of egg-spots on the anal fin.

## Conclusions

### Hypothesis about the genetic basis of the origin and evolution of convergent novel anal fin pigmentation patterns in cichlid fishes

Our findings drive a reassessment with respect to whether convergent pigmentation pattern formation in fish is as the same as that in mammals and birds, whose pigmentation pattern formation is controlled through a few genes *(MC1R* and *Agouti*) (Hoekstra et al. 2006; Uy et al. 2016); or should be viewed with respect to a gene network. Although anal fin pigmentation patterns in cichlid fishes are emerging models, stripe pattern formation in zebrafish can provide some clues. Unlike in mammals, pigment cells in fish are quite diversified, and different pathways are involved in their differentiations (see review from (Parichy 2009)). Stripe pattern formation is driven by the interactions among different pigment cells, and mathematical models have been developed to explain the interactions (Kondo & Miura 2010; Watanabe & Kondo 2015; Kondo 2017). Different pigment cells also have different roles in driving pattern formation in different species (Singh & Nüsslein-Volhard 2015; Patterson et al. 2014). Genes related to endocrine system, such as thyroid hormone, can also affect stripe pattern formation by affecting the development of pigment cells (McMenamin et al. 2014). The mutations of many genes can induce irregular stripe pattern formation in zebrafish, and different genes are involved in pigmentation pattern formation in cichlid fishes, as we mentioned above. These findings reflect that pigmentation pattern formation in fishes is polygenic and involved in different pathways.

Therefore, we propose one hypothesis with respect to a gene network to explain the origin and evolution of convergent novel anal fin pigmentation patterns in cichlid fishes (Figure 6). On one hand, the common genetic basis, comprising pigment cells, signalling pathways and metabolic pathways, is important for the emergence of these novel traits (intrinsic factor, developmental constraint). These genes could be derived from deep homology (Shubin et al. 2009; McCune & Schimenti 2012), i.e. pre-existing pigment cells on the anal fin (new structures can evolve by deploying pre-existing regulatory circuits (Shubin et al. 2009)), or evolve independently. Accelerated evolutionary rates of common genes and pathways help conserve these novelties during evolution (extrinsic factor, selection). The similar effects of the initial mutations affecting the common gene network (e.g., pathways and cascade of changes) inspired the emergence of these convergent novel traits. On the other hand, independently evolved associations with patterning related TFs and signalling pathways are important for the origin and evolution of egg-spots. Theses connections form a relatively independent gene network that frees egg-spots to evolve into diversified phenotypes (different numbers, positions, sizes and colours) (intrinsic factor, developmental constraint). The relatively homogenous anal fin background could also prompt this diversification. Simultaneously, the integration of advantageous genes, especially those related to signalling pathways into the corresponding gene network can conserve egg-spots and prompt its evolution under selection (extrinsic factor, selection). This hypothesis can provide a clue to identify egg-spots as one of the key innovations to the adaptive radiation in cichlid fishes, which was proposed previously based on phylogeny and character reconstruction (Salzburger et al. 2005a, 2007; Salzburger 2009). In this case, although both are novelties, egg-spots can be viewed as “a novelty out of novelty” compared to the blotch. Further thorough study including more samples across the phylogeny in cichlid fishes will be helpful to test this hypothesis. This hypothesis here can further illuminate the mechanism of the origin and evolution of novelties in a broad sense.

**Figure 6.**
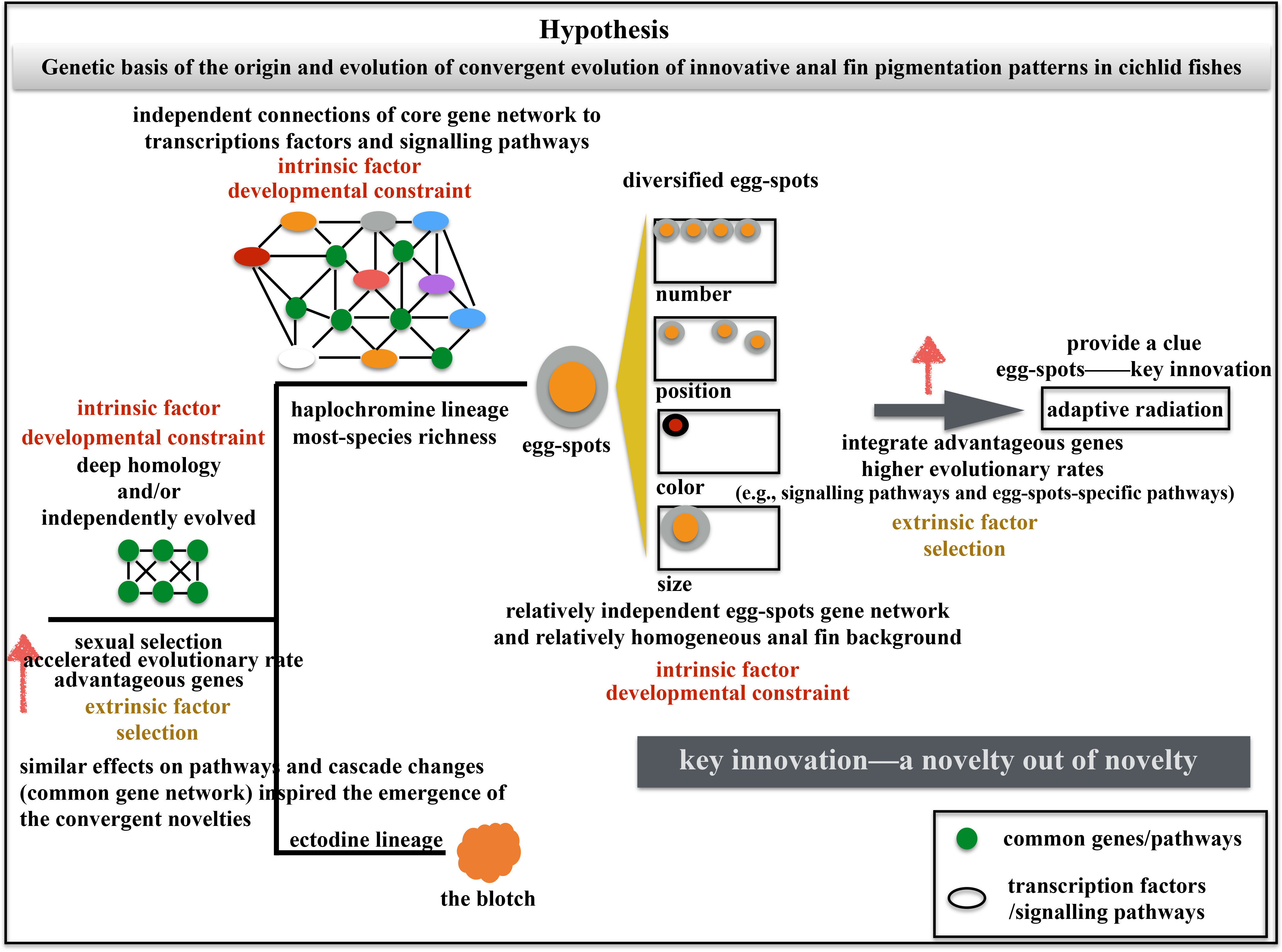
Hypothesis with respect to a gene network to explain the origin and evolution of convergent novel anal fin pigmentation patterns in cichlid fishes. On one hand, the common genetic basis is important for the emergence of these novel traits (intrinsic factor, developmental constraint). These genes could be derived from deep homology or evolve independently. Accelerated evolutionary rates of the common genetic basis help conserve these novelties during evolution (extrinsic factor, selection). Similar effects of the initial mutations affecting the common gene network (pathways and cascade of changes) inspired the emergence of these convergent novelties. On the other hand, independently evolved associations with patterning related TFs and signalling pathways are important for the origin and evolution of egg-spots. Theses connections form a relatively independent gene network that frees egg-spots to evolve into diversified phenotypes (intrinsic factor, developmental constraint). The relatively homogenous anal fin background could also prompt this diversification. Simultaneously, the integration of advantageous genes, especially those related to signalling pathways and egg-spots specific pathways, into the corresponding gene network can conserve egg-spots and prompt its evolution under selection (extrinsic factor, selection). This hypothesis provide a clue to identify egg-spots as one of the key innovations to the adaptive radiation in cichlid fishes (Salzburger et al. 2005a, 2007; Salzburger 2009). In this case, although both are novelties, egg-spots can be viewed as “a novelty out of novelty” compared to the blotch.

## Methods

### Illumina-based RNAseq

Laboratory strains of *C. macrops* were maintained at the University of Basel (Switzerland) under standard conditions (12-h light/12-h dark; 26°C, pH=7). Prior to tissue dissection, the specimens were euthanized with MS 222 (Sigma-Aldrich, USA) following approved procedures (permit nr. 2317 issued by the Cantonal Veterinary Office Veterinary Office). For RNAseq of *C. macrops*, we dissected the anal fins from three adult males and three adult females with similar body sizes. In the male fins, we separated the blotch area from the remaining fin tissue; in the females, which do not possess the blotch, we separated the fin in the same way, i.e., the areas corresponding to the blotch and non-blotch tissue in males resulting in a total of 12 samples for *C. macrops* (Figure 1A). RNA extraction was performed using TRIzol^®^ reagent (Invitrogen, USA). Sample clean up and DNase treatments treatments were performed using the using the RNA clean&Concentrator™-5 (Zymo Research Corporation, USA). RNA quality and quantity was determined using Nanodrop 1000 spectrophotometer (Thermo Scientific, USA) and Bioanalyser 2100 (Agilent Technologies, Germany). Libraries were generated using the Illumina TruSeq RNA Sample Preparation Preparation Kit (low-throughput protocol) according to the manufacturer’s instructions. Per tissue, 330 ng of RNA was subjected to mRNA selection. Pooling and sequencing were performed at the Department of Biosystems Science and Engineering (D-BSSE), University of Basel and ETH-Zurich. Single-end sequencing of these pooled 12 samples was performed in one lane of an Illumina Genome Analyser IIx (maximum read length was 50 bp). The existing raw transcriptomic data for anal fin with egg-spots and without egg-spots of *A. burtoni* was retrieved from Santos et al. (M Emilia Santos et al. 2016).

### Differential gene expression analysis for the blotch and egg-spots

Quality assessment of the sequence reads was conducted using Fastqc 0.10.1 (<u>www.bioinformatics.babraham.ac.uk/projects/</u>). Contaminated Illumina adapter were removed using cutadapt 1.3 (Martin 2011). The reads were subsequently aligned to the Tilapia transcriptome assembly available from Broad Institute (<u>ftp://ftp.ensembl.org/pub/release-81/fasta/oreochromis_niloticus/cdna/</u>. version 0.71). NOVOINDEX (<u>www.novocraft.com/</u>) was used for indexing the reference and NOVOALIGN (<u>www.novocraft.com/</u>) was used for mapping the reads against the reference. The output SAM files of read mapping were subsequently transformed into the BAM format using SAMtools (Li et al. 2009). The count files were concatenated into count tables and analysed using the Bionconductor R package (Gentleman et al. 2004; Huber et al. 2015) and edgeR (Robinson & Smyth 2007; Robinson et al. 2010; Zhou et al. 2013). For paired tissues, we used the individual as the block factor. In the analyses with edgeR, genes that achieved at least one count per million (cpm) in at least three samples were maintained. Differentially expressed transcripts were maintained if the false discovery rate (FDR) was smaller than 0.01.

To obtain blotch candidate genes to the largest extant, the position and sexual effects were controlled (for details see Figure 1A). We did not use the same strategy to find egg-spots candidate genes, since the females of *A. burtoni* also possess egg-spots, although they are less conspicuous than those in males (Theis et al. 2017). Therefore, we corrected the position effect for egg-spots using position genes retrieved from *C. macrops*. Considering the very closely related genetic background in cichlid fishes (Baldo et al. 2011; Brawand et al. 2014) and the conservative fin development constraint (Maxwell & Wilson 2013; Paço & Freitas 2017), the effect of species-specific position genes should be small, and thus, we can get the egg-spots candidate genes to the largest extant. In addition, for finding common genes between the blotch and egg-spots, the comparison itself has already perfectly corrected the position effect considering their different locations on the anal fin (Figure 1).

### Comparative transcriptomics between the blotch and egg-spots

Gene expression clustering was visualized as a heatmap using the *pheatmap* R package v1.0.8 (<u>https://cran.r-project.org/web/packages/pheatmap/index.html</u>). GO annotation of the differential expressed transcripts was conducted with Blast2GO version 2.5.0 (Conesa et al. 2005). BLASTx searches were achieved using BLASTx (threshold: e-6) against zebrafish *(Danio rerio)* protein database and a number of hits of 10. KEGG (Kyoto Encyclopaedia of Genes and Genomes) pathway annotation was conducted with KAAS (KEGG Automatic Annotation Server) http://www.genome.jp/tools/kaas/ (Moriya et al. 2007) using zebrafish as the reference with a threshold e-value of e^−10^. Enrichment analysis of the GO terms was performed using DAVID (<u>https://david.ncifcrf.gov/</u>) with tilapia genome as the background. Protein-protein interaction network analysis was conducted with the string database (<u>http://string-db.org/</u>) with tilapia genome as the reference. The FCNs were extracted and visualized with Cytoscape v3.0 (Shannon et al. 2003). The bootstrap method was used to detect the significance of interaction degrees. Briefly, we randomly extracted the same numbers of the genes as we would like to detect in the network for 10,000 times, and calculated the corresponding average degree every time to get the distribution first. Then, we tested whether the average degrees of the target genes was located outside the 95% confidence interval.

### Evolutionary rates detection of DE transcripts and related pathways

To detect the evolutionary rate of the DE transcripts, we extracted the corresponding transcripts of tilapia retrieved from the Ensembl database (<u>http://www.ensembl.org/index.html</u>) first. These transcripts were used as references after removing stop codon with Biopython (http://biopython.org/wiki/Biopython). The raw reads of transcriptomic data from the other four available cichlid fishes *(Pundamila nyererei, Neolamprologus brichardi, Maylandia zebra* and *A. burtoni*) were retrieved from the NCBI database (https://www.ncbi.nlm.nih.gov/). These data, together with transcriptomic data of egg-spots and the blotch, were mapped to individual transcripts trimmed to the reference using Geneious (Kearse et al. 2012). To retrieve recent duplicated genes, we differentiated the variable single nucleotide polymorphisms deriving from duplicates by mapping these anomalies against the reference. To do this, we concatenated all the transcripts for each species using Geneious followed by visual assessment. Subsequently, we constructed two rounds of consensus sequences through pairwise alignment between the target species and tilapia using MAFFT, which removes the variable single nucleotide polymorphism according to the reference. The resulting concatenated sequences were re-mapped using existing transcriptome data and translated into amino acids followed by visual assessment. Afterwards, individual transcripts were retrieved with Geneious.

To detect the evolutionary rate of these DE transcripts, we aligned the sequences using T_Coffee (Notredame et al. 2000) based on the codon. Subsequently, pairwise Ka/Ks values were calculated using yn00 in Ka/Ks calculator (Wang et al. 2010). To keep the comparison unbiased, we removed the gene pairs when any NA (not applicable or infinite) value occurred (Wang et al. 2011). To determine whether the same pattern also occurred in a lineage-specific manner, we detected the dN/dS values for individual DE transcripts under the free-ratio model within codeml in PAML (Yang 1997, 2007).

## Authors’ contributions

LG designed the experiment, performed the RNAseq library construction and did data analysis. LG constructed the figures and did the analysis of interaction degree of gene networks with the help of CX. Both authors read and approved the manuscript.

## Acknowledgements

We particularly thank Prof. Walter Salzburger for the support. We also thank Dr. Emilia M. Santos for providing the raw transcriptomic data of egg-spots. Many thanks to Chao Zhang, Shengkai Pan, Yongjian Zhang, Lei Luo, Yongsheng Li, Jie Zhang, Tingting Zhou, De Chen and Guang Yang for the discussion and suggestion. We thank Philippe Demougin and Ina Nissen for the assistance with Illumina sequencing. We also thank Attila Rüegg for the assistance in the aquarium room and suggestions. Many thanks to Nicolas Boileau for the assistance in the lab and Brigitte Aeschbach for the laboratory apparatus organization. Pictures were provided by Heinz Büscher, Anya Theis and Adrian Indermau. This work was supported by the PhD grant from University of Basel, Switzerland and Postdoc funding from Southwest University, China. Illumina reads from this study are available at NCBI under the accession number SRP082469. Alignment sequences are available upon request from the corresponding author.

---

**Additional File 1**: Illumina sequencing reads for the ectodine blotch in *C. macrops.*

**Additional File 2**: Evolutionary rates detection of pathways for the blotch and egg-spots.

